# High dose dietary vitamin D allocates surplus calories to muscle and growth instead of fat via modulation of myostatin and leptin signaling

**DOI:** 10.1101/2022.05.19.492715

**Authors:** Caela Long, Zahra Tara, Alex Casella, Julian Mark, Jeffrey D. Roizen

## Abstract

Obesity is the leading proportional cause for diabetes, heart disease and cancer. Obesity occurs because the body stores surplus calories as fat. Fat cells secrete a hormone, leptin, that modulates energy balance at the brain. Changes in fat mass are mirrored by changes in serum leptin. Increases in leptin cause the brain to decrease appetite and increase energy expenditure. However in obesity, leptin sensitivity is decreased which mutes leptin mediated changes in appetite and energy expenditure. We have limited understanding of what controls leptin production by fat or how sensitive the brain is to leptin. Muscle produces a hormone, myostatin, that plays an analogous role to the role that leptin plays in fat. Absent myostatin leads to increased muscle mass and strength. We also do not know what controls myostatin production or sensitivity. Although fat mass and muscle mass are closely linked, the interplay between leptin and myostatin remains unexplored. Vitamin D improves lean mass via what are thought to be primarily trophic effects at the muscle. Here we show that high dose dietary vitamin D preferentially allocates excess calories to muscle and growth instead of storage as fat by decreasing myostatin production and increasing leptin production and sensitivity. That is, high dose vitamin D improves organismal energy sensing. Obesity, aging and other chronic inflammatory diseases are associated with decreased muscle function and mass. Our work provides a physiologic framework for how high-dose vitamin D would be effective in these pathologies to increase allocation of calories to muscle instead of fat and reveals novel interplay between the myostatin and leptin signaling whereby myostatin conveys energy needs to modulate leptin effects on calorie allocation. Furthermore, our work reveals how physiologic seasonal variation in vitamin D may be important in controlling season-specific metabolism and calorie allocation to fat in winter and muscle in summer.

## INTRODUCTION

Vitamin D signaling has been implicated as possibly conveying nutrient status to the brain(1). Vitamin D signaling is mediated primarily by activation of the vitamin D receptor (VDR) to bind DNA and alter transcription(2). VDR knockout mice have a complex failure-to-thrive phenotype with abnormal serum calcium and phosphate, poor growth and nearly-absent white fat(3). Their failure to thrive occurs because they have decreased storage of lipids into white fat(3–6). White fat produces the majority of circulating leptin. Thus, the lack of white fat in VDR knockout mice causes them to have persistently low leptin(3–6). This low leptin causes increased appetite(7).

In addition to nutrient sensing, vitamin D also plays a role in muscle function. Low vitamin D has long been known to cause muscle weakness that is relieved by replenishment of vitamin D. Animal models to date have primarily examined vitamin D signaling and effects through experimental models of deficiency. Specifically, investigators have used approaches focusing on limitation of dietary vitamin D, or conventional or tissue specific knockout of the VDR(8–12). A number of clinical studies, however, suggest that increasing vitamin D within the normal range may have additional beneficial effects on muscle function(13–17).

These functions of vitamin D, regulation of fat metabolism, and regulation of muscle function, have largely been examined separately. However, fat mass and muscle mass are closely linked. Interventions that increase muscle mass increase weight and fat mass. Conversely, in adults when weight loss exceeds 10% of baseline weight, muscle mass loss begins to exceed fat mass loss. Nonetheless, the mechanisms linking fat mass and muscle mass are poorly understood.

Conventional VDR knockout mice have very low serum leptin and muscle specific VDR knockout mice have excess production of myostatin mRNA(3, 9). We wanted to examine the relationship between vitamin D signaling, fat mass, muscle mass, and muscle function. Further, we wanted to examine this relationship not only by comparing normal-to-low dietary vitamin D but also by comparing normal-to-high dietary vitamin D.

To examine the hypothesis that high-dose dietary vitamin D provides muscle function benefit beyond normal-dietary vitamin D, we sought to determine the extent to which increasing vitamin D within the normal range (from 20-30 ng/dL to above 30 ng/dL) improved muscle function in adult wild-type mice. Normal dietary vitamin D increased strength over low-dose vitamin D without altering lean mass. As hypothesized, high dietary vitamin D more dramatically improved strength beyond normal vitamin D. However, high-dose vitamin D also increased lean mass without any change in weight. That is, high-dose vitamin D caused the redistribution of calories from fat to muscle.

Replenishment of vitamin D from low to normal decreased serum myostatin and increased the amount of leptin produced per fat mass. High-dose vitamin D increased sensitivity to leptin without significantly affecting the amount of leptin produced per fat mass. This increased leptin sensitivity did not alter appetite but did increase fat free mass adjusted energy expenditure. High-dose vitamin D also increased linear growth and lean mass proportion of weight. Thus, here we report for the first time that high dose dietary vitamin D preferentially allocates excess calories to muscle and growth instead of storing them as fat by decreasing myostatin signaling and increasing leptin production and sensitivity.

A variety of common pathologies including obesity, aging and chronic inflammatory diseases are associated with decreased muscle function and mass (e.g. sarcopenia). Our work provides a physiologic framework for understanding how high-dose vitamin D would be effective in these pathologies to increase allocation of calories to muscle instead of fat.

Several groups have attempted to use leptin as a treatment for obesity without significant success. The failure of leptin as a treatment for obesity may reflect that leptin signaling in the context of obesity is beyond the dynamic portion of the leptin response curve. Nonetheless, the possibility of increasing leptin signaling by altering leptin sensitivity has largely been neglected. Our work here thus provides initial evidence for the possibility of manipulation of leptin sensitivity to address obesity. Another possible reason for the failure of leptin as a therapy is that it may be one of several checkpoints in energy balance. The loss of muscle mass once weight loss exceeds 10% suggests that another checkpoint in energy balance may be energy needs. Myostatin has been conceptualized as having homeostatic effects on muscle mass. Our work suggests that myostatin may also work to convey energy needs centrally. Thus, our work further suggests that accounting for energy need signaling by inhibiting myostatin pathways could be used to preserve muscle mass in the context of weight loss.

Finally, our work reveals how physiologic seasonal variation in vitamin D may be important in controlling season-specific metabolism and calorie allocation to fat in winter and muscle in summer. Furthermore, these energy sensing and calorie allocation effects may help to explain seasonal variation in human growth, which is concentrated in the Summer and Fall. This is deeply meaningful in several respects. First, from the framing of vitamin D effects, this result provides a context for understanding other vitamin D effects such as improvement of both female and male fertility and possibly seasonal variation in fertility. Second, viewed through the lens of metabolism, this provides a sub-acute signaling pathway to begin to understand metabolism.

## RESULTS

### High dose vitamin D significantly improves grip strength compared to normal vitamin D

To examine calciometabolic-independent effects of vitamin D on muscle, we used the vitamin D receptor knockout rescue diet (VDRKR diet) and varied vitamin D in wild-type (C57BL/6) mice. We used three levels of vitamin D in the diet: 0 IU/kg (low), 2000 IU/kg (normal) and 10,000 IU/kg (high). After 4 weeks these diets allowed us to achieve target serum 25(OH)D of less than 5 ng/mL, between 20-30 ng/mL and above 30 ng/mL.

To determine the extent to which high-D improves muscle function in a dose-responsive fashion above the bottom of the normal range, we measured grip strength as previously described (18). Normal-D significantly improved grip strength over no-D. High-D more dramatically increased grip strength over normal-D (**Fig 1A, \*\*\***p < 0.001 for each by Tukey’s Post-hoc test). *While the improvement in going from low-D to normal-D is expected, the more substantial improvement in going from normal-D to high-D is novel*.

**Fig 1:**
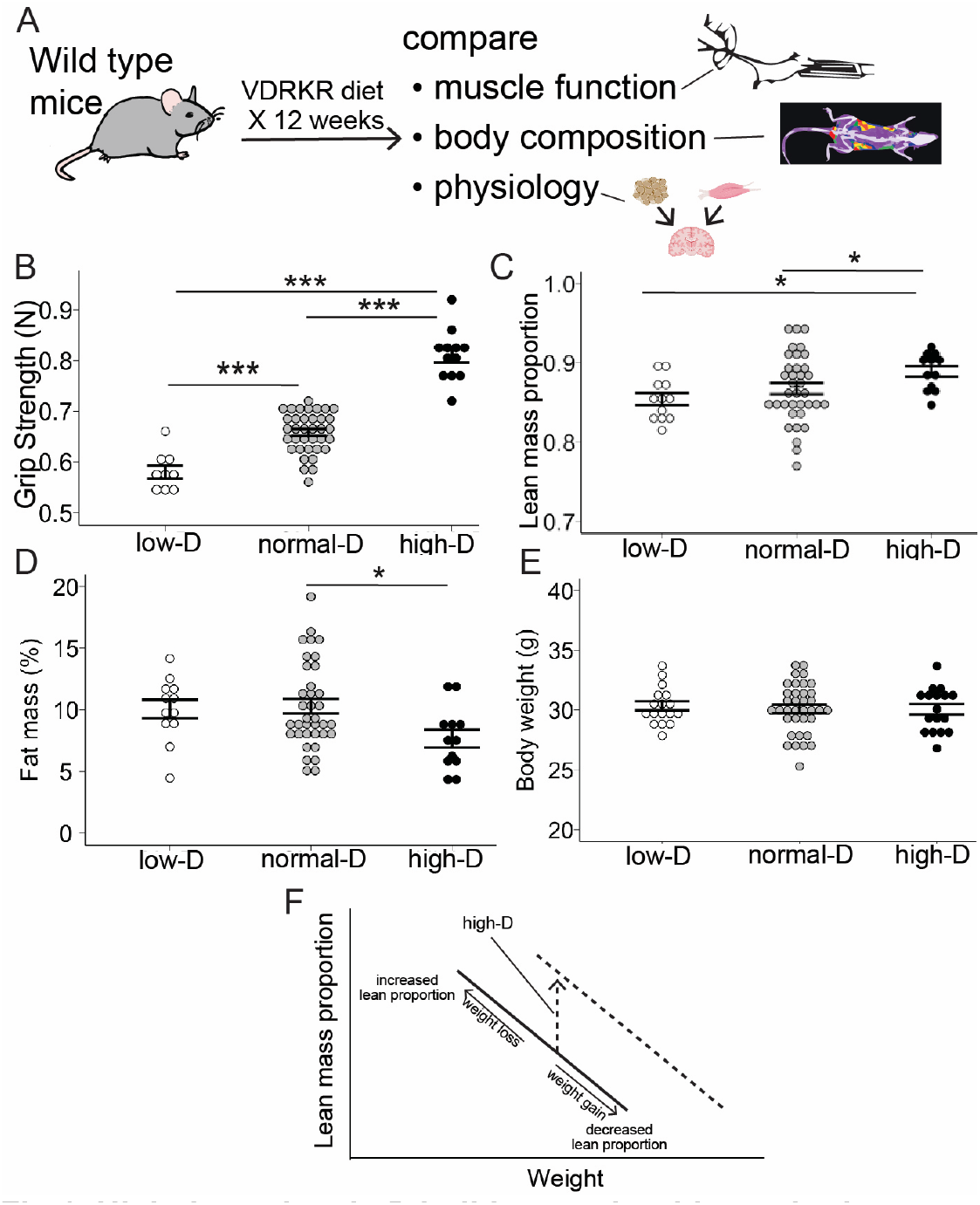
High dose vitamin D builds muscle without altering weight. **A.** Wild-type mice were maintained on a defined vitamin D receptor knockout rescue (VDRKR) diet with each of different doses of vitamin D for 12 weeks and then we assessed the effects of each dose of vitamin D on strength, body composition and physiology. **B**. High-D improves grip strength (*** 0.001>p for each by Tukey post-tests) **C.** High-D improves lean mass (*= p<0.05 for each by Tukey post-tests). **D**. High-D decreases fat mass (*= p<0.05 for each by Tukey post-tests). **E.** High-D decreases fat mass (*= p<0.05 for each by Tukey post-tests). **F.** High-D shifts the curve of the relationship between weight and lean mass. In the linear relationship between lean mass and weight: weight gain decreases the proportion of lean mass and weight loss increases the proportion of lean mass. High-D shifts this curve right such that lean mass proportion increases for the same weight.

### High dose vitamin D increases lean mass, but normal vitamin D does not

To examine the extent to which the high-dose vitamin D stimulated increase in strength reflected changes in muscle mass we measured mouse body composition via NMR as described previously (18). Normal-D did not increase lean mass over low-D but high-D increased lean mass and decreased fat mass relative to normal-D *without altering mouse weight* (**Fig 1B** for lean mass, *: p < 0.05 by ANOVA with Tukey post-tests, **Fig 1C** for fat mass, *: p < 0.05 by ANOVA with Tukey post-tests **Fig 1D** for weight, p > 0.05 by ANOVA). Thus, the high-dose vitamin D stimulated increase in strength is mediated in part by increased calorie allocation to build muscle mass instead of fat mass (schematic in **Fig 1F**).

### Increased vitamin D inhibits myostatin production

Recent work in muscle specific VDR knockout mice as well as in mouse models of vitamin D deficiency reveal that normal vitamin D concentrations facilitate VDR dependent decreases in myostatin mRNA in muscle (12). To examine the possibility that vitamin D dose alters circulating myostatin to regulate the proportion of muscle mass and fat mass, we measured serum myostatin in all three diet groups. Normal D decreased serum myostatin significantly relative to low-D (**Fig 2**, low-D vs normal-D). High-D did not further alter serum myostatin relative to normal-D(**Fig 2**, normal-D vs high-D). This first decrease in myostatin (**Fig 2A**) occurred without any change in fat free mass (**Fig. 1C**) suggesting that normal-D decreased average production of myostatin by muscle mass. The unchanged average myostatin in the high-D diet compared to normal-D occurred in the context of increased average lean mass (**Fig. 1B**), suggesting that high-D further decreased average myostatin production by muscle mass. Overall, these results are consistent with increasing doses of dietary vitamin D inhibiting myostatin production (**Fig 2B**).

**Fig 2:**
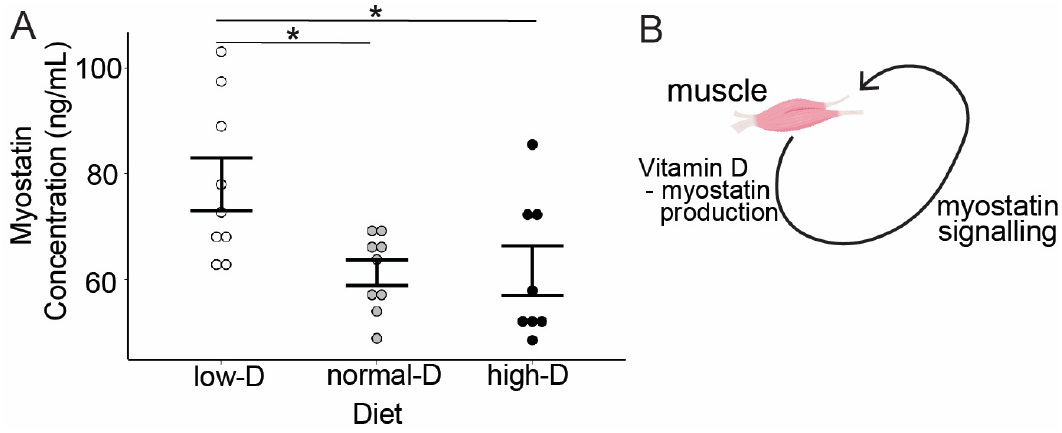
Increasing vitamin D inhibits myostatin production. Wild-type mice were maintained on defined CHOW diets with different doses of vitamin D at sacrifice serum was harvested and myostatin was measured by ELISA. **A.** Normal-D and high-D decrease myostatin concentrations (*= p<0.05, for each ANOVA with Tukey post-tests). **B.** Normal-D and high-D inhibit myostatin production.

### Normalizing vitamin D improves leptin generation per fat mass, but high-dose vitamin D increases leptin sensitivity

Vitamin D receptor deficient mice are hypoleptinemic and are largely deficient in white fat, the type of fat largely responsible for leptin secretion. To examine the possibility that vitamin D dose alters leptin signaling to regulate the proportion of muscle mass and fat mass, we measured serum leptin in all three diet groups. Normal-D increased serum leptin significantly relative to low-D (**Fig 3A**, normal-D vs low-D). High-D decreased leptin significantly relative to normal-D (**Fig 3A**, high-D vs normal-D). To better understand how this pattern related to leptin production and sensitivity, we examined the relationship between serum leptin, fat mass and vitamin D treatment group. Within a vitamin D treatment group, total fat mass determined serum leptin (**Fig 3B,** for no-D r^2^ = 0.97, for normal-D r^2^ = 0.92 and for no-D r^2^ -=0.72). However the slope of this relationship was different between groups (* p<0.05 for Normal-D and High-D vs Low-D by ANOVA with Tukey post-tests). This graph reveals how the relationship between leptin and fat mass shifted significantly and meaningfully between groups (Graph in **Fig 3B**, schematic in **Fig 3C**). The slopes in this graph (**Fig 3B**) are representative of leptin produced per fat mass. Raising vitamin D from low to normal significantly increased the slope of the curve for leptin vs fat mass (*red arrow* in **Fig 3B,** *red lettering* in **Fig 3C**): that is, normalizing vitamin D increased the amount of leptin produced per fat mass (*red arrow* in **Fig 3B**). Raising vitamin D from normal-D to high-D did not further change the slope of this relationship. However, increasing vitamin D from normal-D vs high-D shifted the distribution on this line (*blue arrow* in **Fig 3B,** *blue lettering* in **Fig 3C**). That is, further raising vitamin D from normal to high *did not alter* the slope of the curve for leptin vs fat mass but instead shifted the distribution of fat mass down the curve. This is a shift consistent with high dose vitamin D increasing sensitivity to leptin.

**Fig 3:**
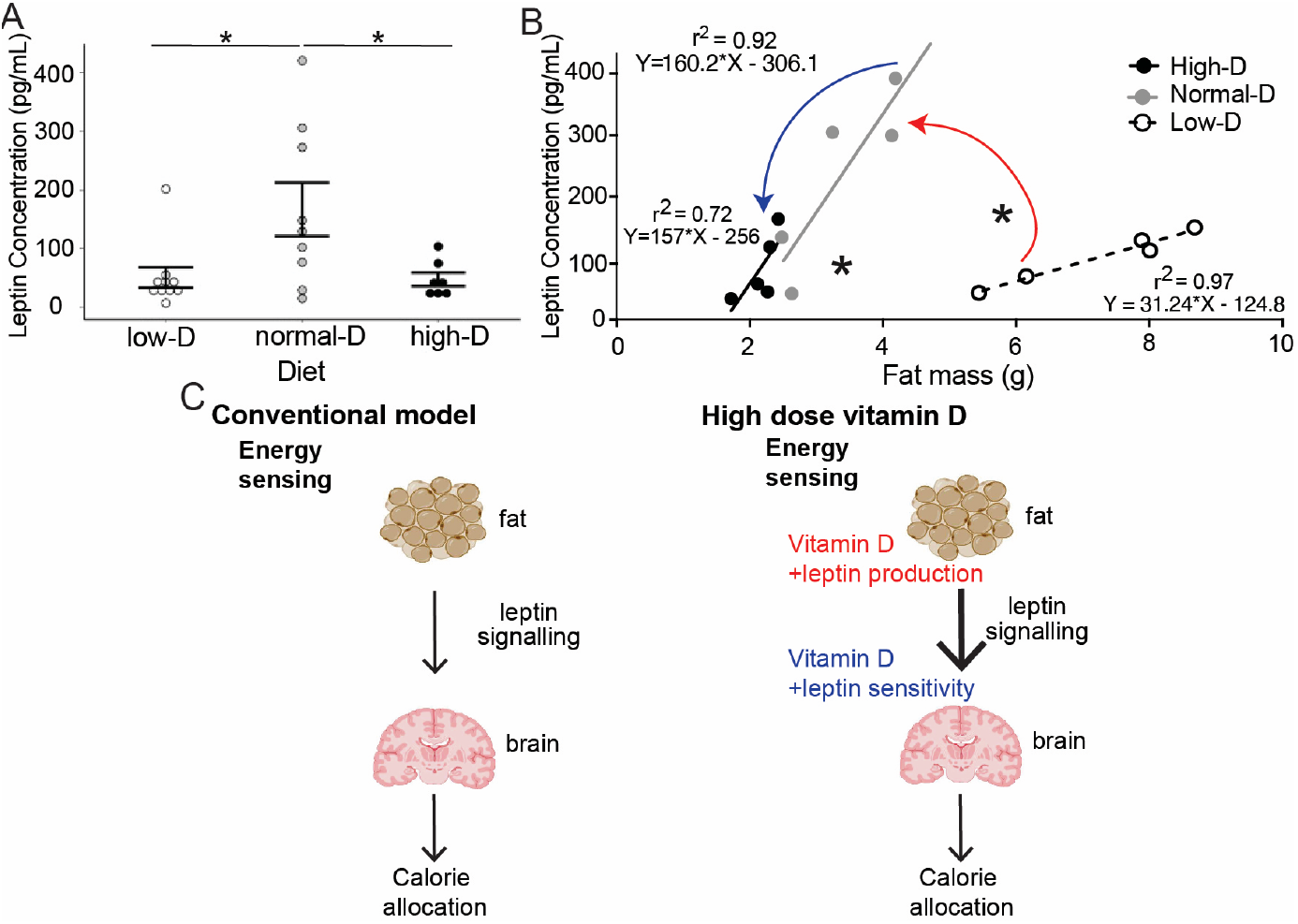
Increasing vitamin D facilitates normal leptin production and improves leptin sensitivity. Wild-type mice were maintained on defined CHOW diets with different doses of vitamin D. At sacrifice leptin was measured by ELISA. **A**: Dietary vitamin D alters serum leptin concentration (*= p<0.05 for each ANOVA with Tukey post-tests). **B**: Dietary vitamin D alters serum leptin production by fat mass; the CHOW low D line is different from both the Reg-D and High-D lines (*= p<0.05 for *= p<0.05 for Normal-D and High-D vs Low-D by ANOVA with Tukey post-tests). **C.** Normalizing vitamin D increased the amount of leptin produced per fat mass (red arrow in **B,** red text in **C**). Raising D from normal to high shifts the distribution of fat masses (blue arrow in **B,** blue text in **C**).

### High dose dietary vitamin D significantly increases energy expenditure without altering activity level or intake

Previously, another group reported that acute paraventricular hypothalamic injection of 1,25D decreases appetite in mice, which was interpreted as increasing leptin sensitivity(1). While some actions of vitamin D may be acute, most actions of vitamin D are mediated by slower transcriptional changes. Thus, it is not clear that this acute response represents a physiologic effect of vitamin D. Leptin has actions on both appetite and energy expenditure, but leptin action is generally thought to alter energy expenditure more than appetite. To further examine dietary vitamin D actions on leptin sensitivity, we measured intake and energy expenditure.

To better understand the apparent increase in leptin sensitivity described above in Figure 3 due to high-dose dietary vitamin D, we investigated the effect of high dose vitamin D on intake and energy expenditure. High dose dietary vitamin D had no effect on mass-adjusted intake (**Fig 4A** and **Fig 4B**, unadjusted intake was also not significantly different between groups). High dose dietary vitamin D had no effect on activity level (data not shown). However, high dose dietary vitamin D significantly increased fat free mass adjusted energy expenditure (**Fig 4C** and **4D**, * p < 0.05 by t-test). Raw-energy expenditure (unadjusted for fat free mass) was also significantly different between groups (**\*** p < 0.05 by t-test, data not shown)) without altering activity level. *Thus high-dose dietary vitamin D signals to differentially allocate calories to muscle instead of fat without altering weight by increasing leptin secretion and leptin sensitivity*.

**Fig 4:**
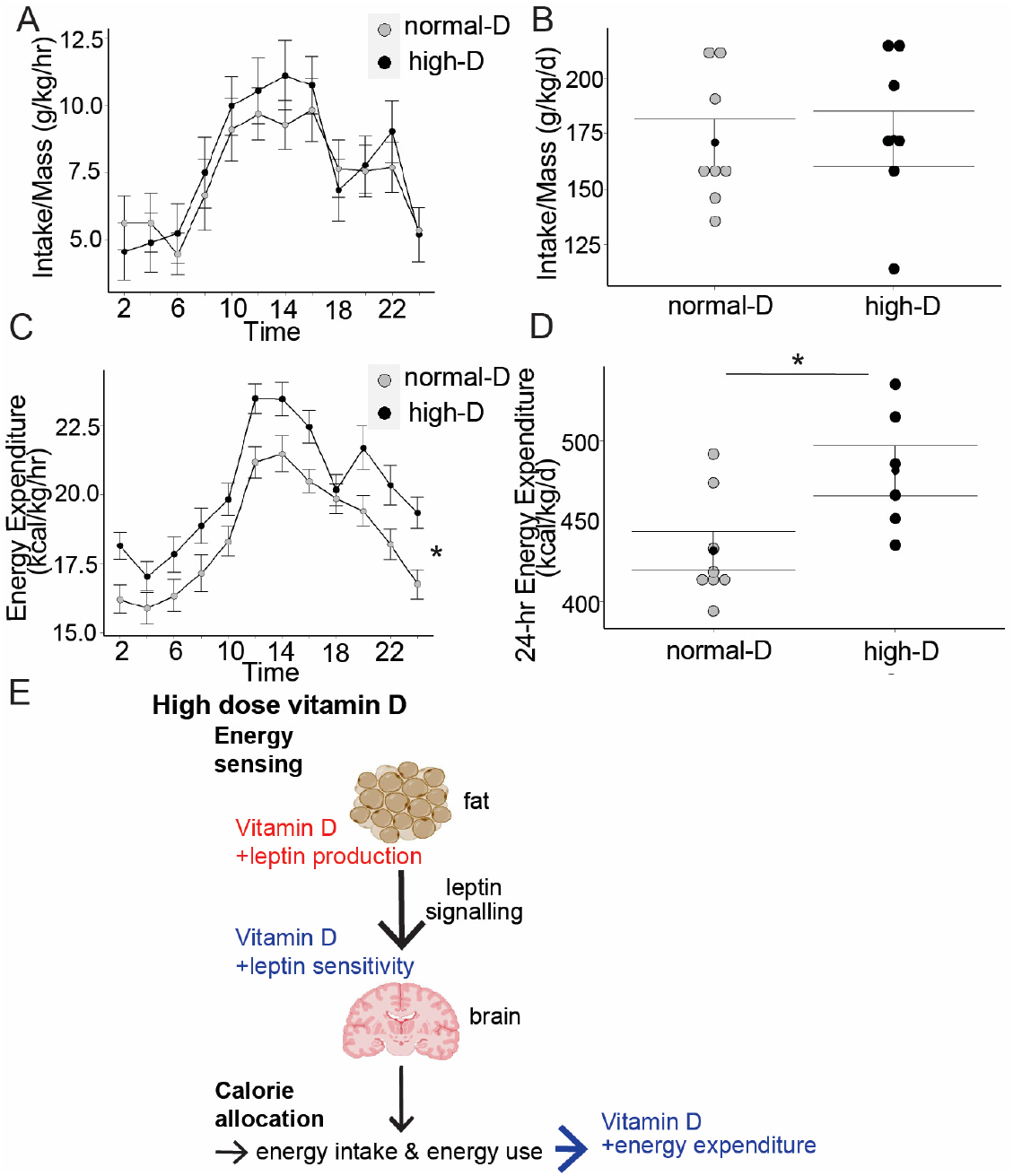
Increasing vitamin D increases energy expenditure without altering intake. Wild-type mice were maintained on defined CHOW diets with different doses of vitamin D. Metabolic cages were used to assay intake, energy expenditure and activity level over one week after acclimatization. **A**, **B**: High dose vitamin D did not alter weight adjusted intake over normal-dose vitamin D. **C**, **D**: High dose vitamin D significantly increased fat-free-mass adjusted 24-energy expenditure relative to normal-dose vitamin D. **E**: High dose vitamin D increases leptin production and sensitivity which raises energy expenditure.

**Fig 5:**
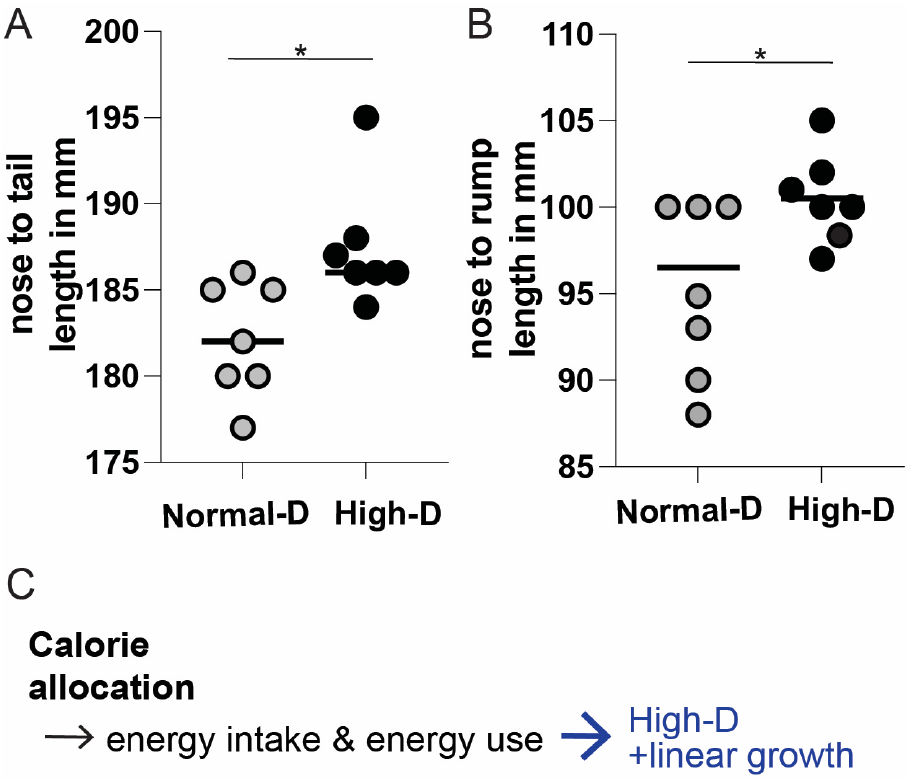
High-dose dietary vitamin D increases length. Wild-type mice were maintained on defined CHOW diets with different doses of vitamin D. **A**. High dose vitamin D increased nose to tail length over normal-dose vitamin D (*= p<0.05 by t-test). **B**. High dose vitamin D increased nose to rump length over normal-dose vitamin D (*= p<0.05 by t-test). **C**. High-dose dietary vitamin D increases linear growth.

### High dose dietary vitamin D significantly increases linear growth

The results to this point suggested to us a model where myostatin was not simply playing a homeostatic role at muscle alone, but was also possibly conveying nutrient needs centrally (**Fig 6** for model). In this context, we expected that the high-D mediated inhibition of myostatin and increased leptin signaling would not only increase energy expenditure but would also facilitate increased growth. To examine this possibility, we measured length in anesthetized mice. As we hypothesized, high-D increased nose-to-tail and nose-to-rump length in mice (p < 0.05 for each by t-test). This result is consistent with vitamin D facilitating increased central energy sensing.

## DISCUSSION

To examine the hypothesis that increasing vitamin D above the normal range provides further benefit beyond replenishment to normal, we sought to determine the extent to which increasing vitamin D within the normal range (from 20-30 ng/dL to above 30 ng/dL) improved muscle function in wildtype mice. Replenishment of vitamin D to normal (20-30 ng/mL) improved strength without increasing muscle mass. However, here we report that high-dose dietary vitamin D further and more dramatically increases strength by increasing muscle mass. Additionally, this increase in muscle mass occurs without a corresponding increase in weight, but instead with a decrease in fat mass. Thus, we propose that high-dose vitamin D causes the redistribution of calories from fat to muscle.

Replenishment of vitamin D to normal decreased myostatin production, but further increases of vitamin D did not alter serum myostatin. Measuring leptin concentrations across dietary groups revealed that replenishment of vitamin D to normal increased the amount of leptin produced per fat mass while high-dose vitamin D increased sensitivity to leptin. Consistent with the large majority of work on leptin actions, this increased leptin sensitivity did not alter appetite, but did increase fat free mass adjusted energy expenditure (**Fig 4**). Thus, here we report for the first time that high dose dietary vitamin D preferentially allocates excess calories to muscle instead of fat by increasing leptin production and sensitivity leading to increased energy expenditure per fat free mass.

These results summarized in **Table 1** at left give rise to the model (**Fig 6**) where raising vitamin D from low to normal increases leptin production by fat, and (on the right in the figure) further raising serum vitamin D levels from normal to high-normal concentrations increases leptin sensitivity (**Fig 6**). Overall, these changes increase allocation of excess calories to muscle mass instead of fat storage.

**Table 1:**
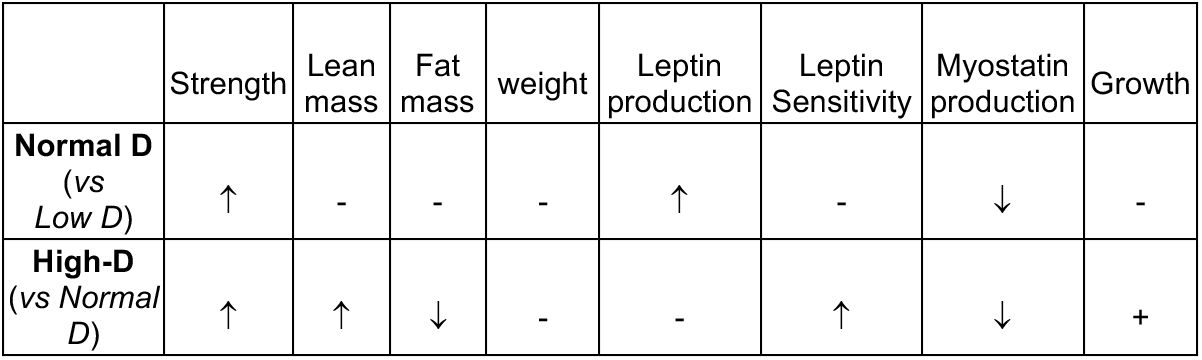
High dose dietary vitamin D preferentially allocates excess calories to muscle and growth instead of fat by increasing leptin production and sensitivity and decreasing myostatin production. Normal-D increases leptin production, decreases myostatin production, leading to increased allocation of excess calories to muscle with increased strength. High-D increases leptin sensitivity and decreases myostatin production, leading to increased allocation of excess calories to muscle with increased strength. as well as increased energy expenditure and linear growth.

A variety of common pathologies including obesity, aging and any disease with chronic inflammation or wasting are associated with decreased muscle function and mass (e.g. sarcopenia). Furthermore, many of these pathologies are associated with low circulating vitamin D because of decreased conversion of calciferols (D2 and D3) to calcidiol (19, 20). Attempts to increase and benefit from an increase in serum Vitamin D have had mixed results in the context of aging, but, recent work in pediatric diseases with chronic inflammation including sickle cell and congenital-HIV identifies significant improvements in muscle function with high-dose vitamin D(21, 22). In many wasting diseases, weight gain is a proxy for increased health. However, careful assessment of how this weight gain is allocated reveals that increases in lean mass specifically improve functional measures of disease (e.g. measures of pulmonary function in cystic fibrosis)(23). Thus, our work provides a physiologic framework for how high-dose vitamin D may be used in these contexts to increase allocation of excess calories to muscle instead of storage as fat.

**Fig 6:**
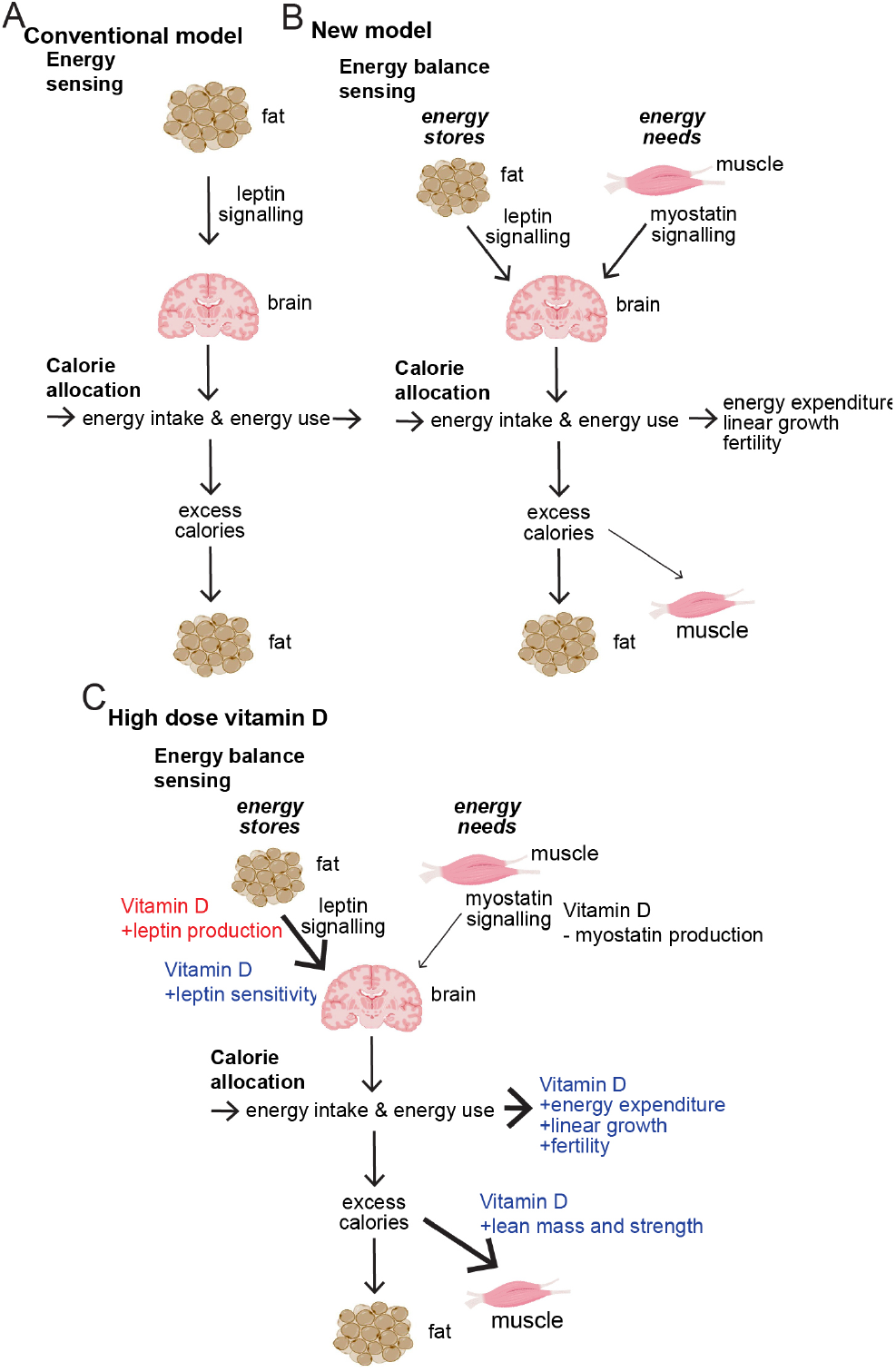
High dose dietary vitamin D preferentially allocates excess calories to muscle and growth instead of fat by increasing leptin production and sensitivity and decreasing myostatin production. In the ***Conventional Model*** (**A** top left) energy sensing is mediated by leptin and by default excess calories are stored as fat. In the ***New Model*** (**B** top right) we propose energy balance is sensed by integrating *energy stores* (conveyed by leptin) and *energy needs* (conveyed by myostatin). In this model we have represented this integration as occurring centrally in the brain but it may occur peripherally. In this model when energy stores exceed needs, some excess calories are used to build muscle and others are used through increased energy expenditure, for growth or to improve fertility. In this model, as pictured at right (in **High dose Vitamin D** (**C**, at bottom)) vitamin D increases leptin production and sensitivity, decreases myostatin production, leading to increased allocation of excess calories to muscle as well as increased energy expenditure and linear growth or improved fertility.

Several groups have studied leptin as a potential therapeutic for obesity. However, aside from the notable exceptional success in leptin-deficient lipodystrophies, leptin has been largely ineffective in combating obesity in clinical trials. The failure of exogenous leptin to effectively treat obesity may reflect that leptin signaling in obesity is already saturated and past the dynamic portion of the leptin response curve. In contrast to the study of exogenous leptin application, exploration of ways to increase leptin signaling or alter leptin sensitivity has been relatively scarce. Our work here not only provides, for the first time, evidence for the possibility that manipulation of leptin sensitivity can address obesity, but it also introduces vitamin D as an initial target for this manipulation. Another possible reason for the failure of exogenous leptin alone as a therapy is that leptin signaling may be only one of several checkpoints in energy balance. The body’s catabolism of muscle once weight loss exceeds 10% suggests that another checkpoint in energy balance may be energy needs. Previously, myostatin has been thought to have homeostatic effects on muscle mass. Our work suggests that myostatin may have an additional role in conveying energy needs centrally. Thus, our work further suggests that addressing energy need signaling by inhibiting myostatin pathways can preserve muscle mass in the context of weight loss.

Finally, our work reveals how physiologic seasonal variation in vitamin D may be important in controlling season-specific metabolism and calorie allocation to fat in winter and muscle and growth in summer. Humans have long been known to grow more in the summer and fall than the spring and winter. This pattern has been ascribed to historic nutrient availability. However, this pattern has persisted in the developed world where nutrient availability is stable throughout the year.

This result is important in several respects. First, from the framing of vitamin D effects, this result provides a context for understanding other vitamin D effects beyond its well-recognized function in calcium homeostasis. Vitamin D improves fertility in females with polycystic ovarian syndrome (PCOS). PCOS is a disease with abnormal or dysfunctional energy balance signaling. Vitamin D also improves fertility in males. In both of these cases, vitamin D may also facilitate season-specific fertility. Second, viewed through the lens of metabolism, this model provides a less acute signaling pathway to begin to understand metabolism. Until now effects of the well understood signaling in energy balance convey nutrient status that changes on a day to day or week to week basis. High-dose dietary vitamin D alters the set points of the leptin and myostatin pathways to convey season specific nutrient availability expectations. With this long term signaling that we have identified, a large feast in the winter would not give rise to increased muscle mass or a growth spurt but would instead be stored as a metabolic insurance against the possibility of scarcity for the remainder of the winter.

## METHODS

### Mouse diets and husbandry

All animal work was reviewed and approved by the Institutional Animal Care and Use Committee of the Children’s Hospital of Philadelphia (Protocol # 0988). To examine calciometabolic independent effects of vitamin D on muscle we used the vitamin D receptor knockout rescue diet (VDRKR diet) and varied vitamin D. We used three levels of vitamin D in the context of defined CHOW VDRKR diets: 0 IU/kg (low – ENVIGO #140078), 2000 IU/kd (normal - ENVIGO #140079) and 10,000 IU/kg (high - ENVIGO #140080). After 4 weeks these diets allowed us to achieve target serum 25(OH)D of less than 5 ng/mL, between 20-30 ng/mL and above 30 ng/mL. The serum calcium, phosphate and PTH were within the normal range in all three groups and were not different between groups. The 1,25(OH)D was undetectable in all three groups. Twelve-week-old male wild-type (WT) C57BL/6J mice (Jackson Laboratories, Bar Harbor, ME) were housed (n = 5 per cage) under a 12:12-h light-dark cycle (light on at 0700) and an ambient temperature of 22°C, and allowed free access to water and diet. Food intake was measured weekly, and body composition was assessed 12 weeks later with nuclear magnetic resonance (NMR) (Echo Medical Systems, Houston, TX).

### Grip strength

Mouse grip strength was measured at the University of Pennsylvania Penn Muscle Institute muscle core with a grip meter (TSE; Bad Hamburg, Germany) as described previously (*18)*. Briefly, mice were trained to grasp a horizontal metal bar while being pulled by their tail and the force was detected by a sensor. Ten measurements were determined for each mouse and averaged.

### Metabolic assays

Body composition of animals was analyzed by NMR at Mouse Phenotyping, Physiology and Metabolism Core, Perelman School of Medicine, University of Pennsylvania. Then, mice were individually housed in the cages, and their metabolic physiology (intake VO2, VCO2 and RER) was monitored by comprehensive laboratory animal monitoring system (CLAMS) at the Mouse Phenotyping, Physiology and Metabolism Core, Perelman School of Medicine, University of Pennsylvania. Myostatin elisa (Biotechne/R & D Systems Myostatin Quantikine ELISA Kit) and leptin elisa (Biotechne/R & D Systems Mouse/Rat Leptin Quantikine ELISA Kit)were performed on serum samples as per manufacturer instruction in technical triplicate.

